# Empirical evidence reveals the phase diagram of patch patterns in Mediterranean drylands

**DOI:** 10.1101/171835

**Authors:** Fernando Meloni, Cristiano R. F. Granzotti, Alexandre S. Martinez

## Abstract

Drylands are ecosystems with limited water resources, often subjected to desertification. Conservation and restoration efforts towards these ecosystems depend on the interplay between ecological functioning and spatial patterns formed by local vegetation. Despite recent advances on the subject, an adequate description of phase transitions between the various vegetated phases remains an open issue. Here, we gather vegetation data of drylands from Southern Spain using satellite images. Our findings support three vegetated phases, separated by two distinct phase transitions, including a continuous phase transition, with new relations between scaling exponents of ecological variables. The phase diagram is obtained without a priori assumption about underlying ecological dynamics. We apply our analysis to a different dryland system in the Western United States and verify a compatible critical behavior, in agreement with the universality hypothesis.

## Introduction

Drylands are arid and semiarid regions covering approximately 40% of the global lands in which more than 2 billions of people live (1). Vegetation in dryland ecosystems occurs in small clusters of plants intercalated by bare soil areas, respectively known as patches and inter-patches. The spatial arrangement of patches (2–4) is important to the ecosystem functionality as a whole, affecting soil stability, chemistry and biology, water runoff, and water absorption (5–8). Small perturbations to the spatial structure of patches may lead to abrupt changes in their ecological structure and functionality. These changes are known as catastrophic shifts. They are linked to desertification processes and may incur economic, environmental and social effects (3, 9–12). Despite recent advances in the area, the consensus on the spatial arrangement of vegetation in drylands remains unclear (13, 14).

Central to this discussion is whether one can sort the various vegetation configurations in Mediterranean dryland (MDL) into groups that share common properties. These groups are the phases of MDL systems. The phase diagram includes transitions between phases and it can provide insights into the circumstances favoring regime shifts, even though the underlying ecological interactions remains poorly known. For instance, the desertification of vegetated regions with decreasing water availability suggests a first-order phase transition (15). In contrast, the large variability observed for patch sizes and shapes observed in MDL empirical data supports discussions concerning scale-free patterns (16–22).

Here, we address the phase diagram of Mediterranean drylands by adopting vegetation patch data as building blocks. Sec. details image collection, from which patch data is extracted. The introduction of a new ecological variable allows us to build the MDL phase diagram. Our findings support three distinct phases separated by two distinct classes of phase transitions, including a continuous one. Critical behavior and critical exponents are further discussed in Sec. . New scaling relations are inferred analytically and, afterward, validated by empirical data. Sec. further characterizes each MDL phase, providing insights on their ecohydrological regime. Our findings emphasize the link between vegetation cover and the spatial arrangement of patches, providing a simple monitoring tool for arid and semi-arid ecosystems.

## Data acquisition and relevant statistics

We consider semi-arid shrublands located in Southern Spain, within the Murcia municipality, which we refer simply as MDL. Vegetation data is acquired from Google Earth images, with a resolution < 40 cm (see Figs. 1), grouped into four image sets. Each set contains non-overlapping and independent images of square plots of lands, hereafter referred as plots, with sides *L*_1_ ≈ 5 m (49 images), *L*_2_ ≈ 10 m (41 images), *L*_3_ ≈ 30 m (40 images), and *L*_4_ ≈ 40 m (55 images), with corresponding areas *A*_*i*_ = *L*_*i*_ × *L*_*i*_. The images are manually processed by color filtering using the ImageJ software (23) as detailed in Ref. (24).

**Fig. 1.**
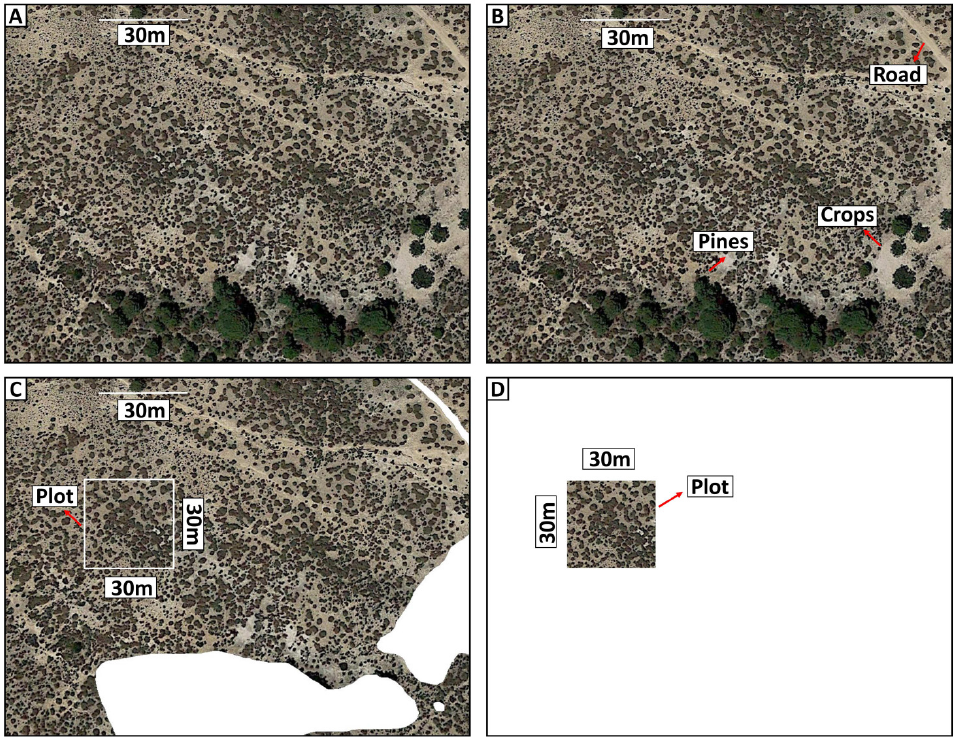
Data acquisition from Google Earth. Plots of land represent the typical shrub vegetation native to the MDL (“Espartales”). To obtain plot samples, random searches are performed through the landscape (view altitude ≈ 200 m), seeking regions containing comprised mostly by target vegetation. (**A**) Raw image acquisition from a selected MDL region. (**B**) Each element in the landscape is identified. (**C**) All undesired elements (crops, roads, buildings, etc) are removed before defining the plot. (**D**) A sample plot image is obtained. Plots are independently sampled, being identified only by their respective geolocation and square area (m^2^).

Plots are independently sampled, representing distinct spatial configurations of patches in MDL, with percentage of vegetation cover area *v* ∈ [0, 1]. The filtered plot images are transformed into binary images, in which pixels can either take value 1 (vegetation) or 0 (bare soil). A cluster of *s* filled pixels surrounded by empty pixels represents a patch of size *s*. For the sake of compatibility with literature data, *s* is measured in m^2^ units after appropriate pixel-to-m^2^ conversion. The variable *N* describes the total number of patches in a given plot, with patch density *D* = *N/A*. Because several patches may co-exist in a plot, we calculate the average size of patches 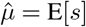 per plot, and its corresponding variance 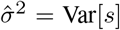. These statistics provide relevant information about the morphology of patches for varying *v*. Conversely, this implies 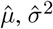 and *N* also depends implicitly on the vegetation cover *v*.

The ecological variables 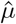 and 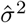 form a good starting point to describe the morphology of MDL vegetation. However, one cannot assume they are suitable order parameters for the complete phase diagram. In part because the pertinent interactions between elements that comprise MDL remain poorly known, thus undermining the identification of symmetries and, hence, order parameters. To solve this issue and construct the phase diagram, it is convenient to introduce a new ecological variable:

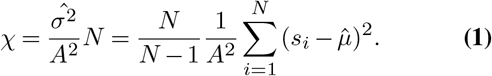

The variable *χ* relates to the total fluctuation for patch sizes in a given plot. Clearly, *χ* is dimensionless and limited *χ* ∈ [0,1]. As a result, data from images with varying plot dimensions can be studied and compared against each other using *χ*.

Fig. 2 depicts the values of *χ* against *v* for plots with size *L*_*k*_(*k* = 1, 2, 3 and 4). *χ* displays three distinctive behaviors separated by phase transitions at *v*_*m*_ = 0.40 and *v*_*c*_ = 0.85. Let us define the phase A for plots with *v* < *v*_*m*_; phase B for *v*_*m*_ < *v* < *v*_*c*_; and phase C for *v* > *v*_*c*_.

**Fig. 2.**
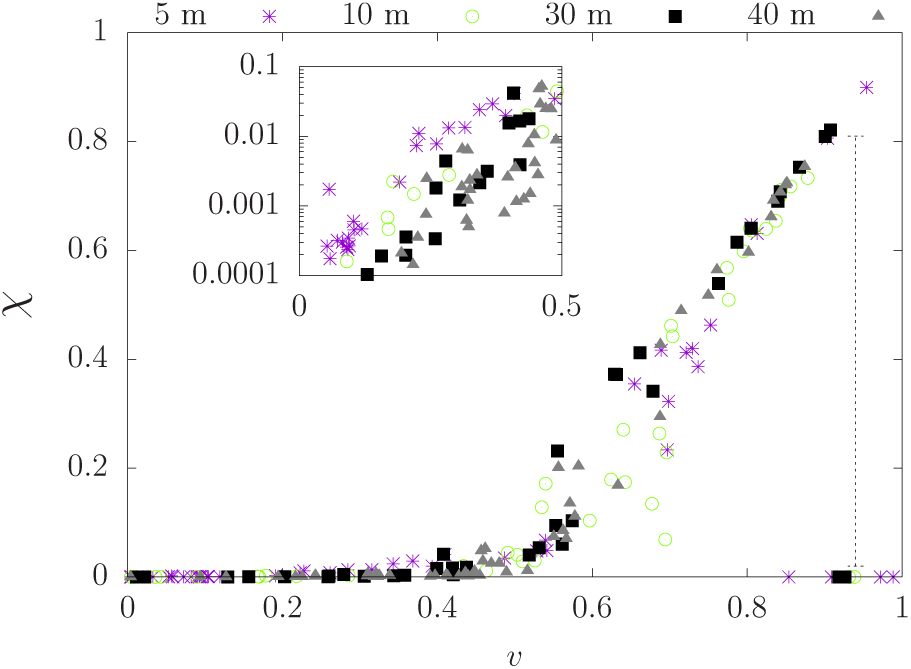
Phase diagram for Mediterranean drylands. The statistic *χ* revels three distinct phases separated by phase transitions at *v_m_* = 0.40 and *v_c_* = 0.85. *χ* increases slowly during phase A, followed by a faster and linear growth rate during phase B. Phase C starts at the discontinuity of *χ*, with vanishing values of *χ*. The vertical dashed line segment indicates the discontinuity at *v_c_*. (inset) *χ* growth is approximately linear in semilog scale, and depends on the plot physical dimensions.

For low vegetation cover *v* < *v*_*m*_ (phase A), patches are small and randomly scattered throughout the plot of land (see Fig. 3). As a result, *χ*, 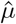, and 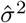 are restricted to small values. In this phase, *χ* grows exponentially albeit very slowly (inset). The parameter of the exponential growth, i.e. the characteristic length scale, depends on plot size. Furthermore, the contact between different patches (coalescence) is unlikely to occur due to the low values of *v*. Hence, increasing *v* in phase A also tend to increase the number of patches *N*.

**Fig. 3.**
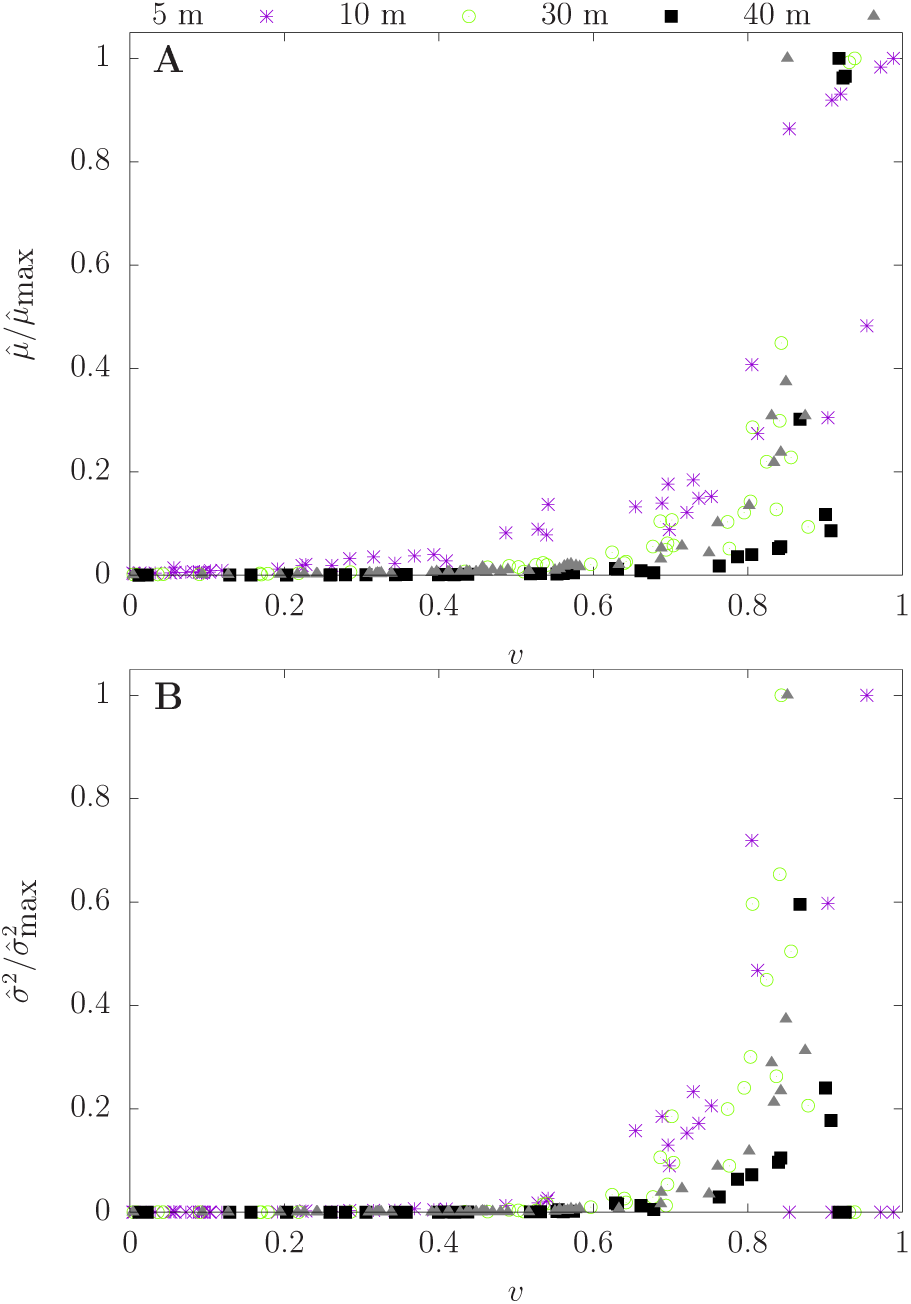
Patch statistics. (**A**) Average patch size normalized by its maximum value taken at four distinct scales, *L*_1_ = 5 m, *L*_2_ = 10 m, *L*_3_ = 30 m, and *L*_4_ = 40m. Around *v*_*c*_ = 0.85, 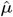 increases rapidly, with vegetation represented by a single large and continuous patch. (**B**) 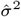 normalized by its maximum value. 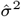 possess vanishing values in most of phase A and phase C. Around *v*_*c*_, it develops a large peak in all four scales.

At *v* = *v*_*m*_, patch coalescence becomes prominent, reducing the number of patches along phase B. At the same time, coalescence between patches also modifies the mean size and variance of patches (see Fig. 3). The transition from the phase with small patches and low fluctuation to a phase with larger patches and larger fluctuations is captured by *χ*. More specifically, for *v* > *v*_*m*_, *χ* grows much faster than in phase A (see Fig. 2), and it can be approximated by a linear function of *v*. This evidence suggests (∂*χ*/∂*v*) exhibits a discontinuity at the phase transition, consistent with a first-order phase transition. In ecological terms, the transition at *v*_*m*_ implies strong dependence on external influences in comparison with the biological interactions. This dependence may be explained as the result of spatial limitations to the increase of *v*, with the coalescence of patches emerging at *v*_*m*_ as an important force affecting patch sizes and vegetation connectivity.

Lastly, *χ* exhibits a discontinuity at *v* = *v*_*c*_, marking the start of the vegetated phase C. Phase C is characterized by one large patch, with average sizes close to the full area of the plot of land, playing the role similar to a giant component in percolation problems. In addition, 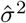 and *χ* vanish in phase C. At the transition *v* = *v*_*c*_, 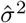 develops a sharp peak in all plot scales (see Fig. 3). We investigate this transition in details in what follows.

## Critical behavior

The normalized values of 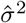 are depicted in Fig. 3. A phase transition takes place around *v*_*c*_ = 0.85, indicated by a prominent peak, suggesting that fluctuations in patch size are important factors for the phase transition. Indeed, around *v*_*c*_, patches in various sorts of shapes and size can be found in MDL plots. The phenomenon occurs in all sets of MDL plots considered in this paper, compatible with the scale invariance that originate collective phenomena. These observations reinforce the hypothesis that a continuous phase transition takes place at *v* = *v*_*c*_.

From a physical perspective, the phase transition separates a non-continuous vegetated phase to a continuous vegetated phase. The statistic 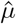 captures the essence of this claim as Fig. 3 depicts, i.e., 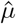 reaches its maximum value for *v* > *v*_*c*_. Hence, 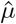 serves as a suitable candidate for order parameter in the continuous phase transition. And because the transition occurs for large vegetation cover, plant-plant interactions and biological effects are more likely to originate the longrange correlations associated with critical phenomena. This in sharp contrast with the phase transition around *v* = *v*_*m*_.

With reasonable evidence of a continuous phase-transition, several inferences can be taken from the theory of critical phenomena (25, 26). In the scaling region, critical exponents *α* and *β* dictate the behavior of the critical phenomena:

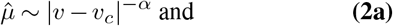

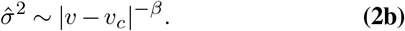

The exponents are estimated from Fig. 3, for each set of plots, using linear fits in log-log scale. The results are *α* = 0.3(9) and *β* = 0.5(2) using plots with size *L*_4_(see Table 1).

**Table 1.** Critical exponents for various sizes of MDL plots of land.

| | $L_1$ | $L_2$ | $L_3$ | $L_4$ |
| --- | --- | --- | --- | --- |
| $\alpha$ | 0.4(9) | 0.4(4) | 0.3(8) | 0.3(9) |
| $\beta$ | 0.6(3) | 0.5(2) | 0.5(3) | 0.5(2) |
| $\Delta$ | 1.1(0) | 1.0(8) | 1.1(1) | 1.1(0) |
| $\gamma$ | 1.4(7) | 1.3(5) | 1.3(2) | 1.3(0) |
| $\rho$ | 1.3(5) | 1.2(3) | 1.2(0) | 1.2(0) |

Although these values match the critical exponent values estimated from numerical models obtained in Ref. (27), Eqs. (2a) and (2b) are subjected to systematic errors due to improper identification of the scaling region (28). Errors become progressively more significant due to lack of plots with large *v* in the data. We mitigate this issue by introducing additional scaling laws that are less sensitive to the scaling region:

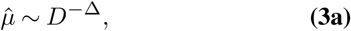

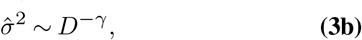

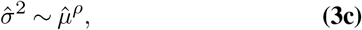

where the scaling exponents Δ, *γ*, and *ρ* are again estimated using linear fits in log-log scales (see Fig. 4). Table 1 summarizes the exponents for all the plot sizes considered in this study. For plots with size *L*_4_, the exponents are Δ = 1.1(0), *γ* = 1.3(0), and *ρ* = 1.2(0).

**Fig. 4.**
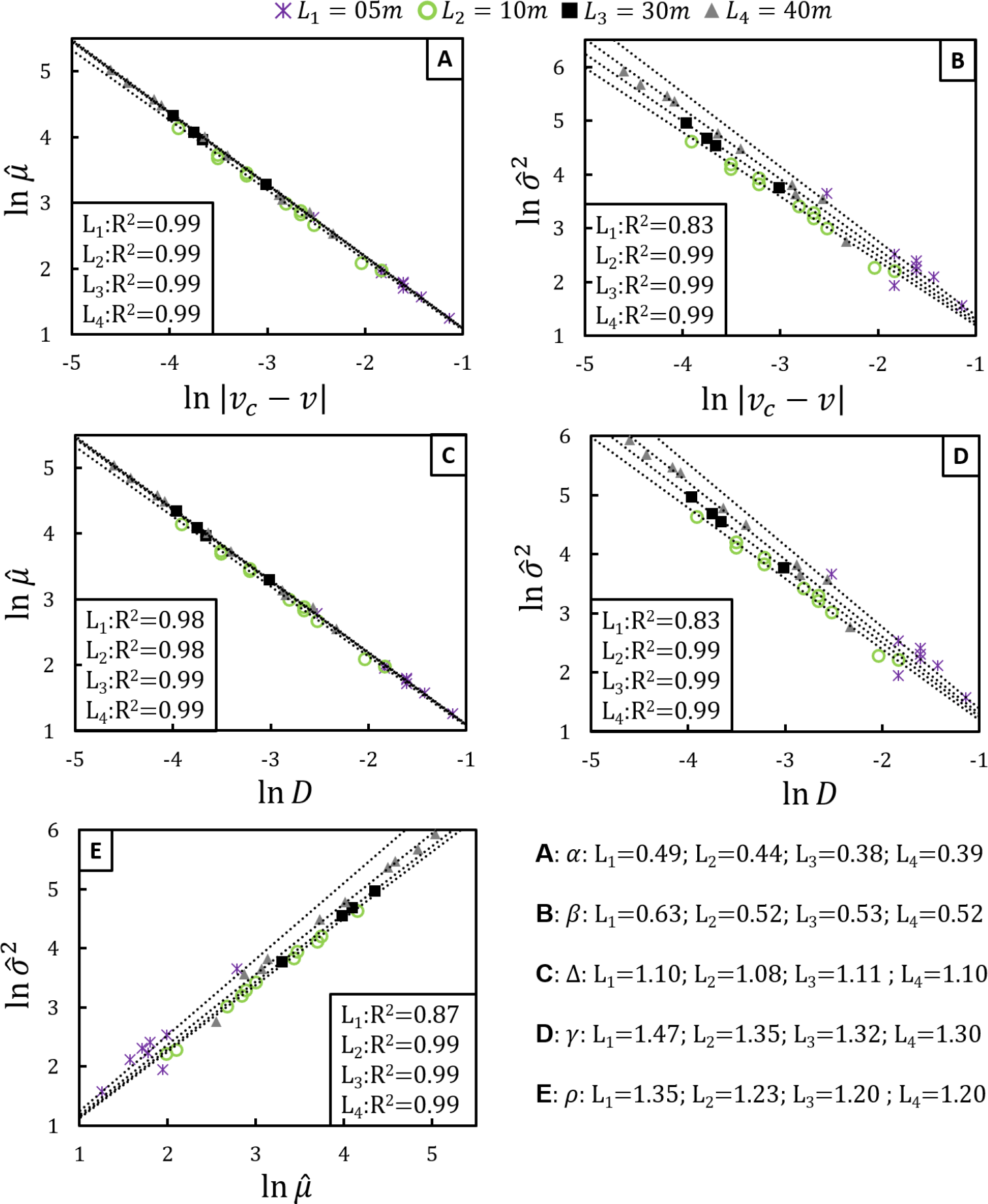
Scaling relations in Mediterranean drylands. Data collected from dry shrublands in south Spain using four distinct plot sizes: *L*_1_ = 5 m, *L*_2_ = 10 m, *L*_3_ = 30 m, and *L*_4_ = 40 m. Scaling relations between: **(A)** average patch size 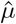 and |*v* −*v*_*c*_|; **(B)** variance 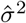 and |*v* −*v*_*c*_|; **(C)** 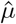 and patch density *D*; **(D)** 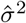 and *D*; **(E)** 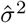 and 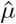.

We further exploit Eqs. (3a-3c) and their low dependence on the scaling region to derive alternative estimates of *α* and *β*. The scaling relations in Eqs. (3a–3c) are consistent if *ρ* = *γ/*Δ. For instance, for plots with size *L*_4_, *γ/*Δ = 1.18 which is compared against *ρ*_data_= 1.20. Next, using Eqs. (2a) and (2b) in Eq. (3c) produces 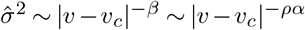. Thus, *β* = *ρα*. Again, using *L*_4_ plots, *β* = *αρ* = 0.47 whereas data fit produces *β*_data_ = 0.52. The results are listed on Table 2.

**Table 2.**
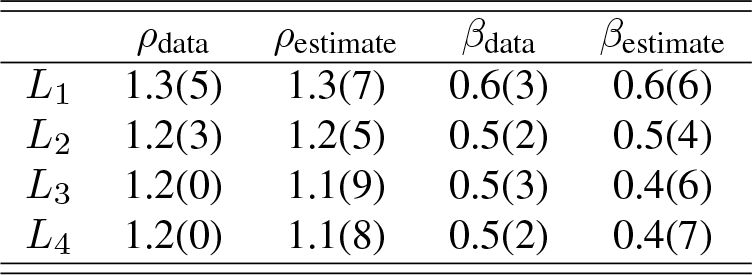
Alternative scaling relations for *ρ* = *γ*/Δ and *β* = *αρ*.

| | $\rho_{\text{data}}$ | $\rho_{\text{estimate}}$ | $\beta_{\text{data}}$ | $\beta_{\text{estimate}}$ |
| --- | --- | --- | --- | --- |
| $L_1$ | 1.3(5) | 1.3(7) | 0.6(3) | 0.6(6) |
| $L_2$ | 1.2(3) | 1.2(5) | 0.5(2) | 0.5(4) |
| $L_3$ | 1.2(0) | 1.1(9) | 0.5(3) | 0.4(6) |
| $L_4$ | 1.2(0) | 1.1(8) | 0.5(2) | 0.4(7) |

## Discussion

Here, we show that MDL supports three distinct vegetation phases (A, B and C) according to the control parameter *v* (see Fig. 5). Phases are separated by a first-order phase transition (*v*_*m*_ ≈ 0.4) followed by a continuous phase transition (*v*_*c*_ ≈ 0.85). Our results contrast with those furnished by traditional models for drylands, in which either a firstor second-order phase transition separating a desertified phase from a completely vegetated one is present (3, 15, 16, 21, 27, 29, 30). In this context, our findings offer evidence and reconcile apparently conflicting views on vegetation patterns in MDL.

**Fig. 5.**
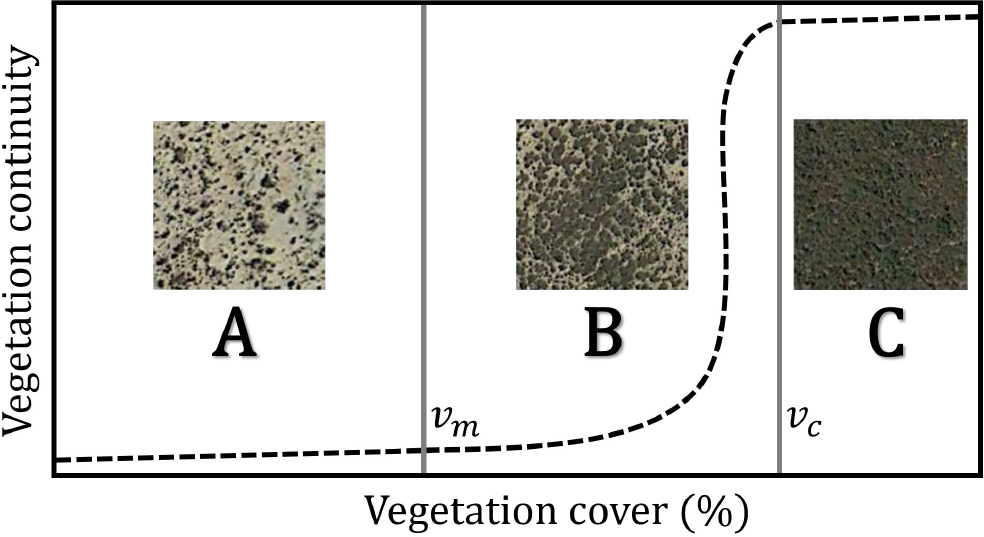
Phase diagram of dry shrublands in south Spain. The figure highlights the distinct spatial arrangement of patches at distinct ranges of vegetation cover *v* according to phases A, B and C. In regime **A**, the geometrical properties of patches show random patterns, with maximum effects of hydric runoff and minimum water infiltration in the soil. A first-order phase transition occurs at *v*_*m*_. In regime **B**, coalescence drives the growth of patches, affecting patch sizes and density in a nontrivial manner. The changes in the spatial arrangement result in a drastic increase in patch connectivity, with consequences for runoff and water infiltration. The coalescence process saturates at the emergence of a percolating vegetation patch, at *v*_*c*_, where a continuous phase transition drives the system from regime **B** to regime **C**. Such a transition is the remarkable consequence of long-range correlations that emerges from biological interactions. In regime **C**, the fully connected vegetation mitigates the effects of runoff and improves water absorption. Terrain features show little relevance, while biological aspects dominate vegetation patterns.

### A. Phases and transitions

Scarce and small patches compromise phase A (*v* < *v*_*m*_), which hinder water infiltration and favor runoff effects (5, 6, 31, 32). Due to their smallness, patches are randomly scattered across the landscape, producing short-range correlations and a characteristic scale. This observation is supported by previous studies in the same region using patch size distribution functions (24, 33). Therefore, facilitation effects are less important in phase A, and terrain influences dominate the vegetation growth (new colonizations and increases at the periphery of already established patches).

As *v* increases towards *v*_*m*_, one also expects the number of patches and their corresponding average size to increase. However, with increasing sizes, nearby patches are more likely to coalesce, reducing the total number of patches. The first-order phase transition is mainly governed by the terrain and climate conditions, which affect the *v*_*m*_ threshold. Hence, *v*_*m*_ is scale-dependent and mainly affected by nonbiological local influences. The changes in scale in drylands have been previously announced in (3). Among their results, they verify numerically that patch size probability distribution function also develops distinct characteristic scales as the vegetation cover increases. The change in characteristic scale at *v*_*m*_ is in agreement with a first-order phase transition.

Phase B occurs within the range *v*_*m*_ < *v* < *v*_*c*_. Unlike the other two phases, the spatial arrangement of patches in phase B lacks a simple characterization as reported by Aguiar and Salas (34). The Authors recognize two phases, low and high cover, for several worldwide drylands with evident differences in spatial patterns and ecological interactions. They describe the low-cover phase as we did for phase A, but they are unable to find a simple description of the high-cover phase. In phase B, the total number of patches decreases, and patches become larger in size. This is a consequence of coalescence, which also increases the variability of geometrical forms and shapes and hinders a simple geometric description of spatial patterns in MDL. The constant competition for space during vegetation growth produces a net increase in the patch variance, culminating in the critical transition at *v*_*c*_.

Our findings concerning the nature of the transition at *v*_*c*_ agree with previous studies (16, 17, 21, 35, 36), but differ in the threshold value *v*_*c*_ proposed in the Ref. (37). For instance, Xu et al. (22) considered several drylands and observed an increase in the skewness of patch size distribution up to *v* ≈ 0.7. However, the Authors were unable to assess larger values of *v*, so that they could not detect the critical phase transition. Here, we consider smaller plots than used in previous studies because it allows a better assessment of the entire *v* range.

In phase C, the vegetation reaches a continuous configuration, prone to water infiltration and minimally affected by runoff. In general, this combination mitigates desertification threats. Moreover, the large levels of vegetation cover also lessen the influence of terrain compared to that of plant-plant interactions as the main regulator of vegetation patterns.

### B. Universality

Bearing in mind the critical phase transition, we highlight the discovery of the scaling relations between the variables *D*, 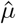, and 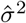. The equalities *ρ* = *γ*/Δ and *β* = *ρα* relate the critical exponents *ρ*, Δ and *γ* to *α* and *β*. This is the expected outcome in the classical theory of critical phenomena, and, as such, grants us confidence in the validity of the critical hypothesis in MDL.

The experimental observation of a critical phase transition is no minor feat. The reason behind this claim lies in another crucial concept of critical phenomena: universality classes (38, 39). This concept asserts that distinct systems exhibit the critical behavior provided they share the same critical exponents and symmetries. In the ecological context, however, the term “distinct systems” concerns distinct drylands, but subject to similar environmental conditions. For drylands with similar conditions and plant-plant interactions, one expects similar critical exponents, i.e., that the various worldwide drylands would be grouped into a reduced number of universality classes.

Recently, Berdugo *et al*. (40) provided empirical evidence that supports the universality hypothesis in drylands. The authors evaluated 115 dryland worldwide, looking for relations between aridity and system functionality. The authors sorted the dryland samples into two groups: high aridity/low functionality associated with a lognormal patch size distribution; high functionality/low aridity environments associated with power-law patch size distributions.

These advances inspired us to extend our analysis drylands located in Utah, US (36 plots). In Utah drylands, patches are constituted by trees associated with shrubs, differing from their counterpart located in Spain. Nevertheless, Fig. 6 depicts that normalized variances from MDL (30 m) and the US (50 m) collapse into a single curve, suggesting a common critical behavior. Furthermore, the exponents found for US drylands are *α*_US_ ≈ 0.42 and *β*_US_ ≈ 0.46 (Supplement-1). As critical exponents are not influenced by terrain details and environmental conditions, this evidence suggests that both regions might share similar interaction rules. Therefore, this test supports our hypothesis of universality classes as a way to sort the various worldwide drylands. The critical transition in both drylands, US and MDL, occurs near the same coverage value, which is an unexpected result because the threshold is not a universal feature.

**Fig. 6.**
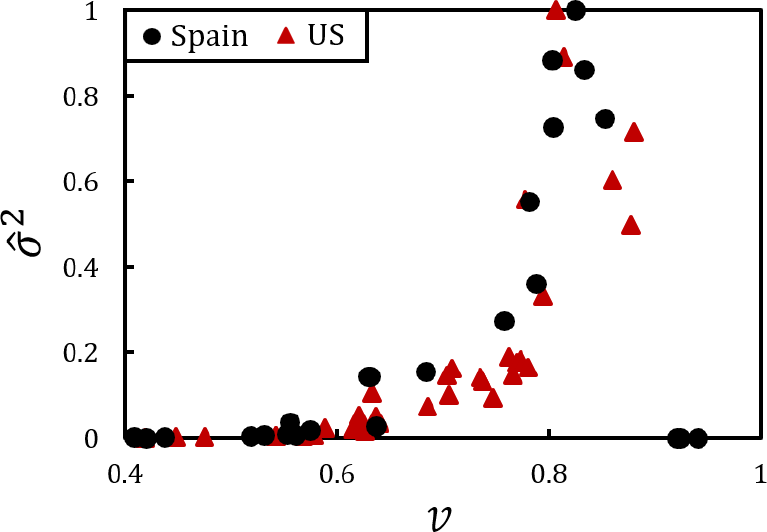
Phase transitions in Mediterranean and Utah drylands. Data from MDL with plots *L*_3_ = 30 m (black full circles), while US data (dark red) uses *L* = 50 m. Despite the inherent ecological differences between both vegetations, their respective normalized variances collapse into a single curve. The values of critical exponents calculated for plots from US *α*_US_ = 0.4(2); *β*_US_ = 0.4(6) are compatible with values calculated for plots from Spain *α* = 0.3(8) and *β* = 0.5(0), respectively.

## Conclusions

We report three vegetation regimes (phases A, B and C) for Mediterranean drylands via patch statistics, using the vegetation cover *v* as control parameter. Phases A and B are separated by a first-order phase transition that takes place at *v*_*m*_ = 0.40. The continuous phase transition between phases B and C occurs near *v*_*c*_ = 0.85. The critical exponents estimates in the largest scale *L*_4_= 40 m are *α* = 0.38 ± 0.02, *β* = 0.5 ± 0.03, Δ = 1.10 ± 0.01, *γ* = 1.30 ± 0.04, and *ρ* = 1.20 ± 0.04. New relations *ρ* = *γ*/Δ and *β* = *αρ* have been analytically determined and validated against empirical data. We also provide new evidence supporting the hypothesis of universality classes in drylands, which can offer insights on the symmetries and inner workings of plant-plant interactions in drylands.

## Ackowledgements

We are grateful for LS dos Santos’ comments during the manuscript preparation. FM thanks S. Bautista for the hospitality during the first stages of the research. ASM holds grants from CNPq 307948/2014-5, GMN thanks CAPES 88887.136416/2017-00, and CRFG acknowledges support from CAPES. This work was supported by the São Paulo Research Foundation (FAPESP) Grant No. 2013/06196-4 and Grant No. 2014/00631-3.

