## Supplementary material for "Empirical evidence reveals the phase diagram of patch patterns in Mediterranean drylands": Supp

### Supplementary Information: Empirical evidence reveals the phase diagram of vegetation patterns in Mediterranean drylands

Fernando Meloni,* Gilberto Medeiros Nakamura, Cristiano Roberto Fabri Granzotti, and Alexandre Souto Martinez†

*Faculdade de Filosofia, Ciências e Letras de Ribeirão Preto, Universidade de São Paulo, Avenida dos Bandeirantes, 3900, 14.040-901, Ribeirão Preto, SP, Brazil*

(Dated: April 8, 2019)

*

† Instituto Nacional de Ciência e Tecnologia em Sistemas Complexos, Rio de Janeiro, RJ, Brazil

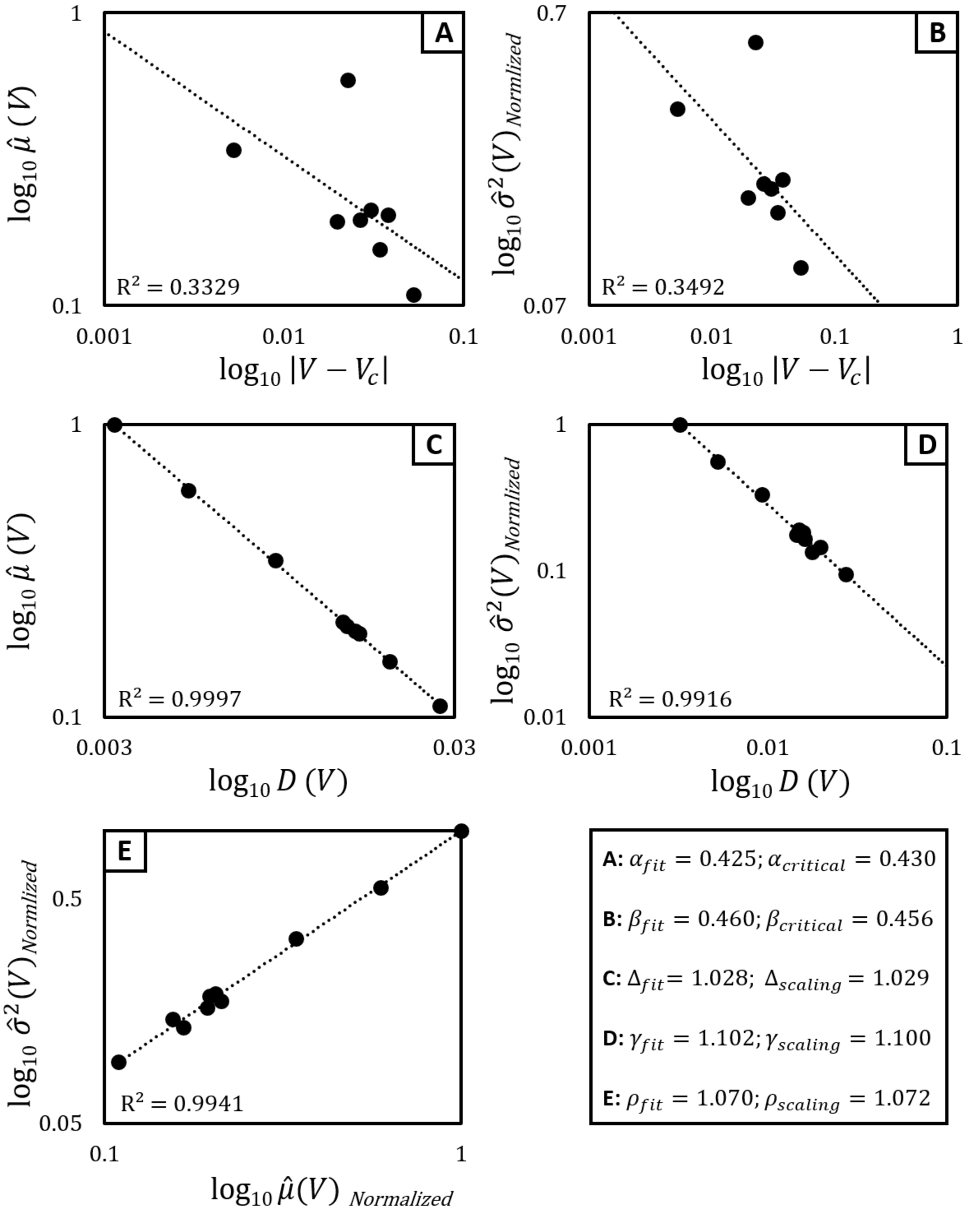


Figure 1. Critical and scaling exponents calculated for a preliminary test applied in plots of drylands from Utah, US. The region shows trees growing together shrubs and, therefore, a different vegetation physiognomy that found in Mediterranean drylands. Despite these differences, the critical exponents are compatible and the scaling relations are maintained.The analysis considered only plots vegetation cover ($V$) around the critical transition ($V_c \approx 0.8$). $\hat{\mu}(V)$ and $\hat{\sigma}^2(V)$: average patch size and respective variance; $D(V)$: density of patches; $\alpha$ and $\beta$: critical exponents; $\rho$, $\gamma$ and $\Delta$: scaling exponents. Exponents values indicated by $fit$ are obtained by the respective angular coefficient of linear fit (log-log scale). Exponent values indicated by *scaling* are obtained by the scaling relations $\beta = \rho\alpha$ and $\rho = \gamma/\Delta$.
